# Whole genome sequence analysis of *Salmonella* Typhi isolated in Thailand before and after the introduction of a national immunization program

**DOI:** 10.1101/076422

**Authors:** Zoe A. Dyson, Duy Pham Thanh, Ladaporn Bodhidatta, Carl Jeffries Mason, Apichai Srijan, Maia A. Rabaa, Phat Voong Vinh, Tuyen Ha Thanh, Guy E. Thwaites, Stephen Baker, Kathryn E. Holt

**Affiliations:** Centre for Systems Genomics, University of Melbourne, Parkville, Victoria 3052, Australia; Department of Biochemistry and Molecular Biology, Bio21 Molecular Science and Biotechnology Institute, University of Melbourne, Parkville, Victoria 3010, Australia; The Hospital for Tropical Diseases, Wellcome Trust Major Overseas Programme, Oxford University Clinical Research Unit, Ho Chi Minh City, Vietnam; Department of Enteric Diseases, Armed Forces Research Institute of Medical Sciences, Bangkok, Thailand; Centre for Tropical Medicine and Global Health, Oxford University, Oxford, United Kingdom; The London School of Hygiene and Tropical Medicine, London, United Kingdom

## Abstract

Vaccines against *Salmonella* Typhi, the causative agent of typhoid fever, are commonly used by travellers, however, there are few examples of national immunization programs in endemic areas. There is therefore a paucity of data on the impact of typhoid immunization programs on localised populations of *S*. Typhi. Here we have used whole genome sequencing (WGS) to characterise 44 historical bacterial isolates collected before and after a national typhoid immunization program that was implemented in Thailand in 1977 in response to a large outbreak; the program was highly effective in reducing typhoid case numbers. Thai isolates were highly diverse, including 10 distinct phylogenetic lineages or genotypes. Novel prophage and plasmids were also detected, including examples that were previously only reported in *Shigella sonnei* and *Escherichia coli*. The majority of *S*. Typhi genotypes observed prior to the immunization program were not observed following it. Post-vaccine era isolates were more closely related to *S*. Typhi isolated from neighbouring countries than to earlier Thai isolates, providing no evidence for the local persistence of endemic *S*. Typhi following the national immunization program. Rather, later cases of typhoid appeared to be caused by the occasional importation of common genotypes from neighbouring Vietnam, Laos, and Cambodia. These data show the value of WGS in understanding the impacts of vaccination on pathogen populations and provide support for the proposal that large-scale typhoid immunization programs in endemic areas could result in lasting local disease elimination, although larger prospective studies are needed to test this directly.

**Author Summary:** Typhoid fever is a systemic infection caused by the bacterium *Salmonella* Typhi. Typhoid fever is associated with inadequate hygiene in low-income settings and a lack of sanitation infrastructure. A sustained outbreak of typhoid fever occurred in Thailand in the 1970s, which peaked in 1975-1976. In response to this typhoid fever outbreak the government of Thailand initiated an immunization program, which resulted in a dramatic reduction in the number of typhoid cases in Thailand. To better understand the population of *S*. Typhi circulating in Thailand at this time, as well as the impact of the immunization program on the pathogen population, we sequenced the genomes of 44 *S*. Typhi obtained from hospitals in Thailand before and after the immunization program. The genome sequences showed that isolates of *S*. Typhi bacteria isolated from post-immunization era typhoid cases were likely imported from neighbouring countries, rather than strains that have persisted in Thailand throughout the immunization period. Our work provides the first historical insights into *S*. Typhi in Thailand during the 1970s, and provides a model for the impact of immunization on *S*. Typhi populations.

## Introduction

*Salmonella enterica* subspecies *enterica* serovar Typhi (*S*. Typhi) is a human restricted bacterial pathogen and the etiological agent of typhoid fever. *S*. Typhi is transmitted faeco-orally and can establish asymptomatic carriage in a small subset of an exposed population (1). Recent estimates (2–4) place the global burden of typhoid fever at 25-30 million cases annually, of which 200,000 are associated with deaths. Typhoid fever occurs most commonly in industrialising countries, specifically in locations with limited sanitation and related infrastructure (5); children and young adults are among the most vulnerable populations in these settings (6–8). Antimicrobial therapy together with water sanitation and hygiene (WASH) interventions are the major mechanisms by which typhoid fever is controlled (9, 10). However, none of these approaches are optimal and resistance against antimicrobials has become increasingly common in *S*. Typhi since the 1970s (11–13). A number of typhoid vaccines are licenced for use (14–18), however, they are not widely used as a public health tools in endemic areas, with the exception of controlling severe outbreaks such as those following natural disasters (19–22).

A sustained typhoid fever outbreak occurred in Thailand in the 1970s. A sharp increase in cases was observed in 1973-1974, which finally peaked in 1975-1976. In response, the government of Thailand established a national typhoid immunization program, which represented the first programmatic use of a typhoid vaccine in the country (23). The immunization program targeted over 5 million school aged children (7-12 years) each year in Bangkok between 1977 and 1987 (80% of the eligible population). Thus, Thai school children were eligible to receive a single locally produced heat/phenol-inactivated subcutaneous dose of 2.5 × 10^8^ *S*. Typhi organisms annually (14, 23), before the program was halted in the early 1990s because of high rates of adverse reactions caused by the vaccine (22). To our knowledge this is the only such programmatic use of a vaccine for controlling Typhoid fever in children in Thailand. Data from four teaching hospitals in Bangkok showed a 93% reduction in blood culture confirmed infections with *S*. Typhi between 1976 (n=2,000) and 1985 (n=132) (14, 23). Notably, no significant decline was observed in isolation rates of *Salmonella* Paratyphi A (*S*. Paratyphi A), a *Salmonella* serovar distinct from *S*. Typhi that causes a clinical syndrome indistinguishable from typhoid fever, but for which *S*. Typhi vaccines provide little or no cross-protection (14). This observation suggests that the reduction in *S*. Typhi infections was not attributable to improvements in infrastructure and hygiene practices only (5, 14, 20, 23). While the inactivated S. Typhi vaccine was found to be highly efficacious (22, 23), it is no longer used as a consequence of being overly reactogenic (14, 16, 23, 24). A Vi capsular polysaccharide vaccine (15) and live-attenuated oral vaccine of strain Ty21a (16) have since replaced this vaccine for travellers to endemic locations (5, 21, 24).

The typhoid immunization program in Thailand provided a unique opportunity to investigate the impact of immunization on *S*. Typhi populations circulating within an endemic area. Here we present an analysis of a historical collection of 44 *S*. Typhi isolates obtained from patients in Thailand between 1973 and 1992 (before and during the immunization program). As *S*. Typhi populations demonstrate little genetic diversity, we used whole genome sequencing (WGS) to characterise these isolates, and core genome phylogenetic approaches to compare the historic isolates from Thailand to a recently published global *S*. Typhi genomic framework (4).

## Materials and methods

### Ethics statement

This is a retrospective study of bacterial isolates unlinked to patient information and was not subject to IRB approval.

### Bacterial isolation and antimicrobial susceptibility testing

Forty-four *S*. Typhi isolated from patients with suspected typhoid fever attending hospitals in Bangkok, Nonthaburi, Loi, and Srakaew, in Thailand between 1973 and 1992 were available for genome sequencing in this study (**Fig 1** and **Table S1**). At the time of original isolation, bacterial cultures were transferred on nutrient agar slants to the department of Enteric Diseases, Armed Forces Research Institute of Medical Sciences (AFRIMS), Bangkok, Thailand for identification and antimicrobial susceptibility testing. At AFRIMS, bacterial isolates were subcultured on Hektoen Enteric agar (HE) and identification was performed by biochemical testing on Kligler iron agar slants, tryptone broth for indole, lysine decarboxylase medium, ornithine decarboxylase medium, urease test, mannitol and motility media (Becker Dickenson, Thailand). Serological agglutination was performed using *Salmonella* O antisera and *Salmonella* Vi antiserum (Difco, USA). Bacterial strains were stored frozen at -70°C in 10% skimmed milk or lyophilised in 10% skimmed milk; lyophilized ampoules were stored at 2-8°C. Prior to DNA extraction for sequencing, lyophilized bacteria was rehydrated with trypticase soy broth, inoculated on McConkey agar and incubated at 37°C for 18-24 hours. If bacteria was stored frozen in skimmed milk, organisms were inoculated directly onto McConkey agar after thawing and then incubated at 37°C for 18-24 hours.

**Figure 1.**
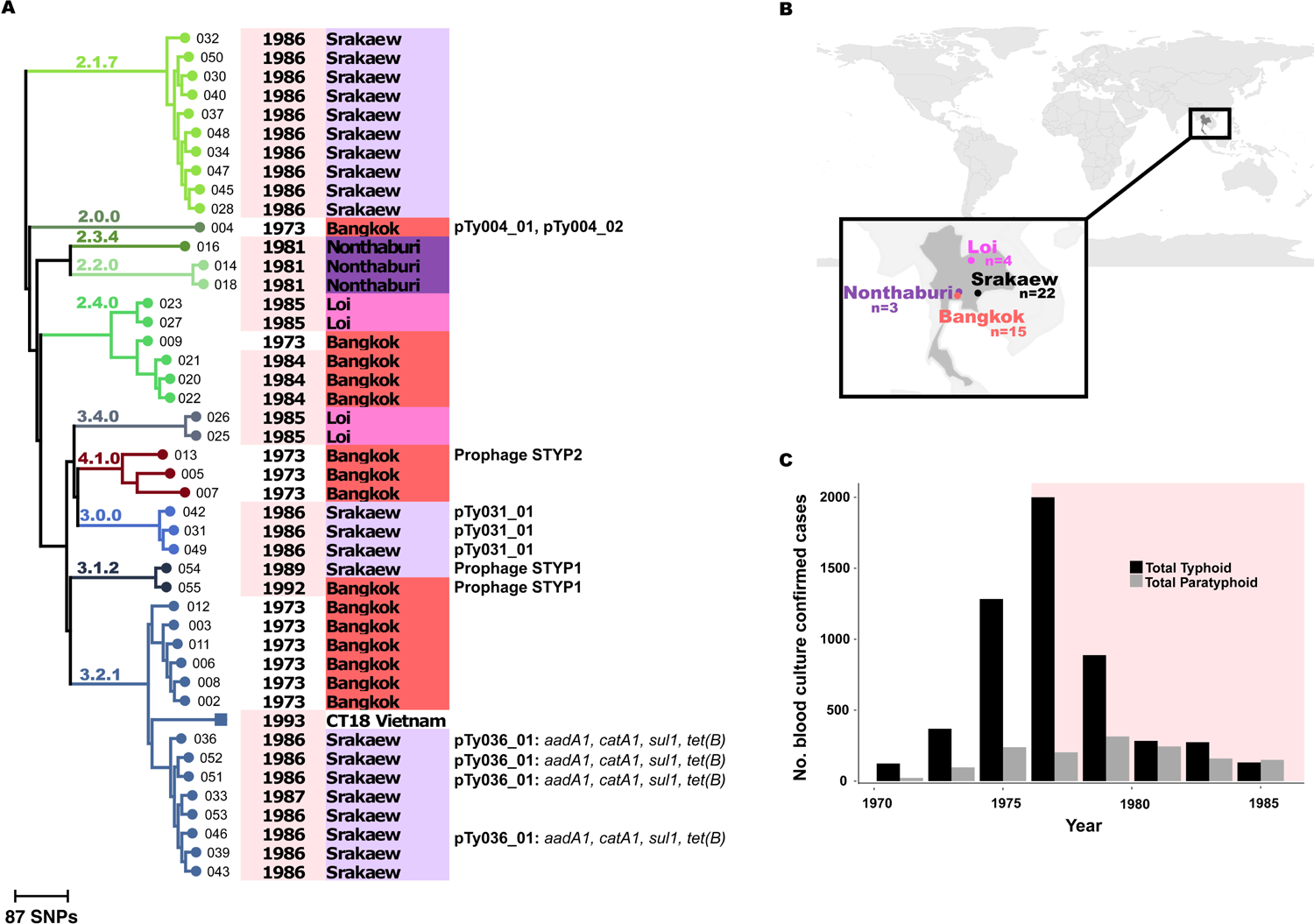
Genomic analysis of Thai *S*. Typhi.

(A) Maximum likelihood phylogenetic tree (outgroup rooted). Strains are labelled with their three digit name code, year of isolation (pink shading indicates post-vaccine isolates); source location (shaded by city, as indicated in panel B); and plasmid content (any antibiotic resistance genes are indicated in italics). Branch lengths are indicative of the number of SNPs. (B) Locations from which S. Typhi were isolated in Thailand. (C) Total number of positive blood cultures of S. Typhi (black) and Paratyphi A (grey) between 1970 and 1985; immunization period is indicated in pink; reproduced using data from reference (14).

Antimicrobial susceptibility testing against ampicillin, chloramphenicol, cephalothin, gentamicin, kanamycin, neomycin, sulfisoxazole, trimethoprim/sulfamethoxazole, and tetracycline was performed by disk diffusion according to Clinical and Laboratory Standards Institute (CLSI) (25–28).

### Genome sequencing and SNP analysis

Genomic DNA from the 44 *S*. Typhi from Thailand was extracted using the Wizard Genomic DNA Extraction Kit (Promega, Wisconsin, USA). Two μg of genomic DNA was subjected to indexed WGS on an Illumina Hiseq 2000 platform at the Wellcome Trust Sanger Institute, to generate 100 bp paired-end reads. For analysis of SNPs, paired end Illumina reads were mapped to the reference sequence of *S*. Typhi CT18 (accession no: AL513382) (29) using the RedDog (v1.4) mapping pipeline, available at https://github.com/katholt/reddog. RedDog uses Bowtie (v2.2.3) (30) to map reads to the reference sequence, then high quality SNPs called with quality scores above 30 are extracted from the alignments using SAMtools (v0.1.19) (31). SNPs were filtered to exclude those with less than 5 reads mapped or with greater than 2.5 times the average read depth (representing putative repeated sequences), or with ambiguous base calls. For each SNP that passed these criteria in any one isolate, consensus base calls for the SNP locus were extracted from all genomes (ambiguous base calls and those with phred quality scores less than 20 were treated as unknowns and represented with a gap character). SNPs with confident homozygous allele calls (i.e. phred score >20) in >95% of the *S*. Typhi genomes (representing a ‘soft’ core genome of common *S*. Typhi sequences) were concatenated to produce an alignment of alleles at 45,893 variant sites. The resultant allele calls for 68 of these SNPs were used to assign isolates to previously defined lineages according to an extended *S*. Typhi genotyping framework (32) code available at https://github.com/katholt/genotyphi). SNPs called in phage regions, repetitive sequences (354 kb; ~7.4% of bases in the CT18 reference chromosome, as defined previously (33) or recombinant regions (~180kb; <4% of the CT18 reference chromosome, identified using Gubbins (v1.4.4) (34)) were excluded, resulting in a final set of 1,850 SNPs identified in an alignment length of 4,275,037 bp for the 44 isolates. SNP alleles from Paratyphi A strain 12601 (35) were also included as an outgroup to root the tree. For global context, raw read data (4) were also subjected to genotyping analysis and those isolates sharing the genotypes that were observed in the Thai collection (n=340; details in **Table S2)** were subjected to the same SNP analyses, resulting in a final set of 9,700 SNPs for a total of 386 isolates.

### Phylogenetic and SNP analysis

Maximum likelihood (ML) phylogenetic trees **(Figs 1-2)** were constructed using the 1,850 and 9,700 bp SNP alignments, respectively, using RAxML (v 8.1.23) (36) with a generalized time-reversible model and a gamma distribution to model site specific recombination (GTR+Γ substitution model; GTRGAMMA in RAxML), with Felsenstein correction for ascertainment bias. Support for ML phylogenies was assessed via 100 bootstrap pseudoanalyses of the alignments. For the larger tree containing global isolates, clades containing only isolates from only a single country were collapsed manually in R using the drop.tip () function in the *ape* package (37). Subtrees were extracted for each subclade, which are therefore each rooted by the other subclades. Pairwise SNP distances between isolates were calculated from the SNP alignments using the dist.gene() function in the *ape* package for R (37).

### Accessory genome analysis

Acquired antimicrobial resistance (AMR) genes were detected, and their precise alleles determined, by mapping to the ARG-Annot database (38) of known AMR genes using SRST2 v0.1.5 (39). Plasmid replicon sequences were identified using SRST2 to screen reads for replicons in the PlasmidFinder database (40, 41). Raw read data was assembled *de novo* with SPAdes (v 3.5.0) (42) and circular contigs were identified visually and extracted using the assembly graph viewer Bandage (v0.7.0) (43). These putative plasmid sequences were annotated using Prokka (v1.10) (44) followed by manual curation. Where IncHI1 plasmid replicons were identified using SRST2, and their presence confirmed by visual inspection of the assembly graphs, IncHI1 plasmid MLST (pMLST) sequence types were determined using SRST2 (13, 45, 46). Where resistance genes were detected from short read data, Bandage was used to inspect their location in the corresponding *de novo* assembly graph in order to determine whether they were encoded in the bacterial chromosome or on a plasmid. Assembled contigs were concatenated and putative prophage genomes were identified with the PHAge Search Tool (PHAST) (47), and their novelty determined by BLASTN analysis against the GenBank database. Pairwise alignments between novel and known prophage sequences were visualised using the *genoPlotR* package for R (48).

**Nucleotide sequence and sequence read data accession numbers**

Raw sequence data have been submitted to the European Nucleotide Archive (ENA) under project PRJEB5281; individual sample accession numbers are listed in **Tables S1 and S2.** Assembled phage and protein sequences were deposited in GenBank, accession numbers are listed in **Table 1**.

**Table 1.**
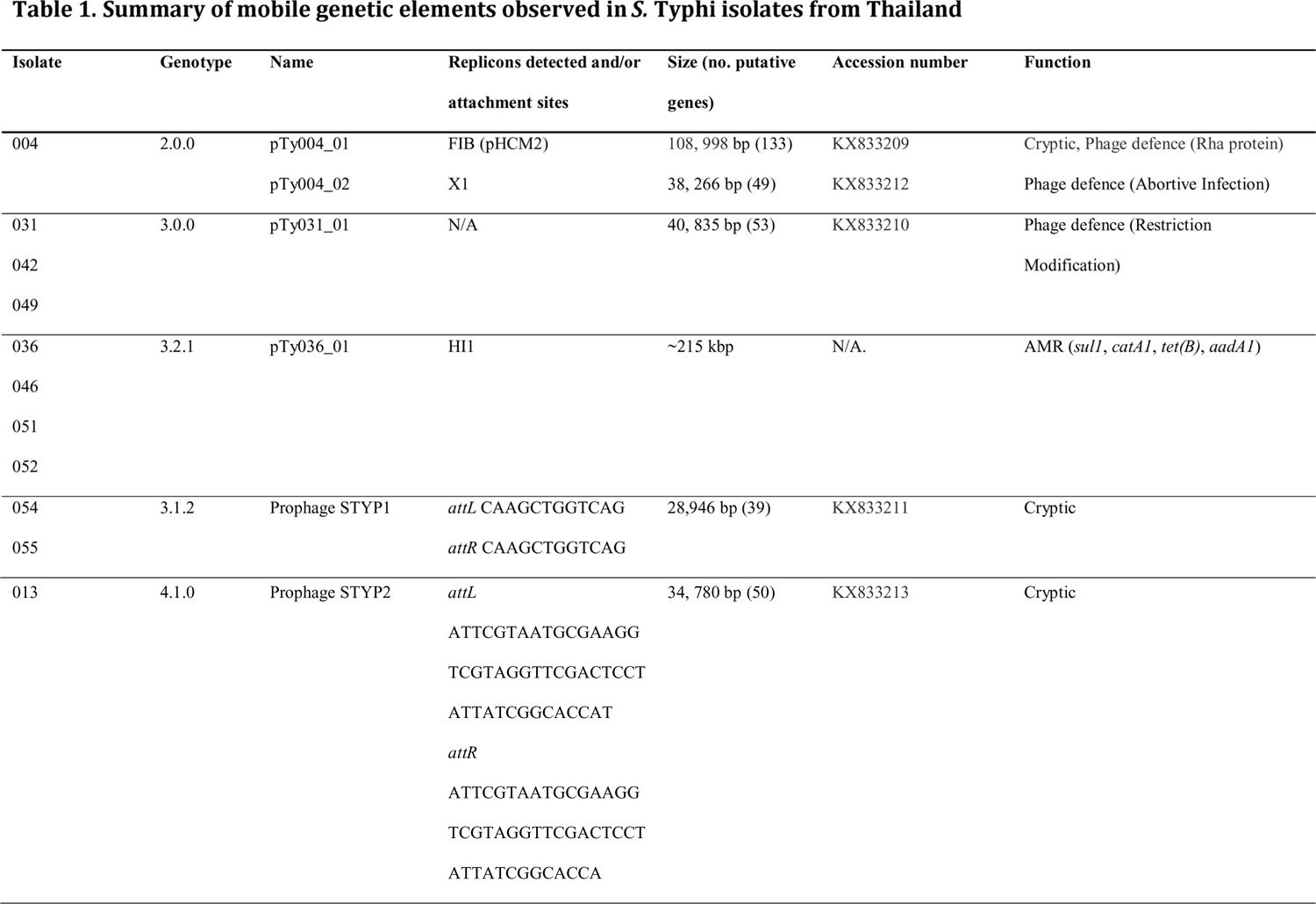
Summary of mobile genetic elements observed in ***S.*** Typhi isolates from Thailand

## Results

### The population structure of S. Typhi in Thailand

All 44 *S*. Typhi isolates collected between 1973 and 1992 were subjected to WGS and SNP analysis. Genome-wide SNPs were used to construct a ML phylogeny and isolates were assigned to previously defined genotypes (32) using a subset of SNPs (see **Methods).** These analyses subdivided the population into ten distinct genotypes, each corresponding to a specific lineage in the ML phylogeny **(Fig 1).** Genotype 3.2.1 (which includes the reference genome CT18, isolated from Vietnam in 1993 (29)) was the most common (n=14, 32%), followed by genotype 2.1.7 (n=10, 23%). Genotypes 2.0 (n=1, 2%) and 4.1 (n=3, 7%) were observed only in 1973 (pre-vaccine period). Genotypes 2.1.7 (n=10, 23%), 2.3.4 (n=1, 2%), 3.4.0 (n=2, 5%), 3.0.0 (n=3, 7%), 3.1.2 (n=2, 5%), were observed only after 1981 (post-vaccine period). Each of these post-immunization genotypes was from a single location and time period **(Fig 1)**, consistent with short-term localised transmission. The only exceptions were the two *S*. Typhi 3.1.2 isolates, that were from Srakaew in 1989 and Bangkok in 1992 and separated by just 4 SNPs. Genotypes 3.2.1 and 2.4.0 were observed amongst both pre-and post-vaccine isolates.

### Thai S. Typhi in the context of a global genomic framework

Based on the Thai *S*. Typhi genotyping results we hypothesised that the post-immunization typhoid infections in Thailand resulted from occasional re-introduction of *S*. Typhi from outside the country, as opposed to long-term persistence of *S*. Typhi lineages within Thailand. To explore this possibility, and to provide a global context for our analysis, we examined 1,832 *S*. Typhi genomes from a recently published global collection that included isolates from 63 countries (4). Genome-wide SNP-based ML trees for each of these genotypes, showing the relationships between Thai and global isolates, are shown in **Fig 2.** In general, post-vaccine Thai isolates were closely related to recent isolates sourced from neighbouring countries including Vietnam, Laos and Cambodia **(Fig 2)**, consistent with regional endemic circulation. In contrast, most pre-vaccine isolates had no close neighbours in the global collection, particularly 2.0.0 strains **(Fig 2A)**, suggesting they may have been Thailand-specific lineages that have died out following the vaccine program. The *S*. Typhi genomes in the global collection were mainly isolated 2-3 decades after the Thai isolates as we did not have access to contemporaneous isolates from these countries that could identify specific transfer events. However, all but three of the post-vaccine Thai isolates shared shorter SNP distances with isolates from neighbouring countries than they did with pre-vaccination Thai isolates (see **Fig 3)**, consistent with these cases being caused by occasional re-introduction of genotypes circulating in the region. Notably, Thai *S*. Typhi 3.2.1 that were isolated in 1986-7 clustered separately from the 1973 pre-vaccine isolates (≥60 SNPs apart), but closely with isolates from Vietnam and Cambodia (differing by as few as 7 SNPs; **Fig 2H).** Post-vaccine Thai *S*. Typhi 2.4 formed two distinct groups that were not consistent with direct descendance from earlier isolates **(Fig 2E).** These data are therefore consistent with transfer of *S*. Typhi into Thailand from neighbouring countries during the post-immunization program era, although the long-term circulation of ancestral populations in Thailand remains an unlikely alternative explanation.

**Figure 2.**
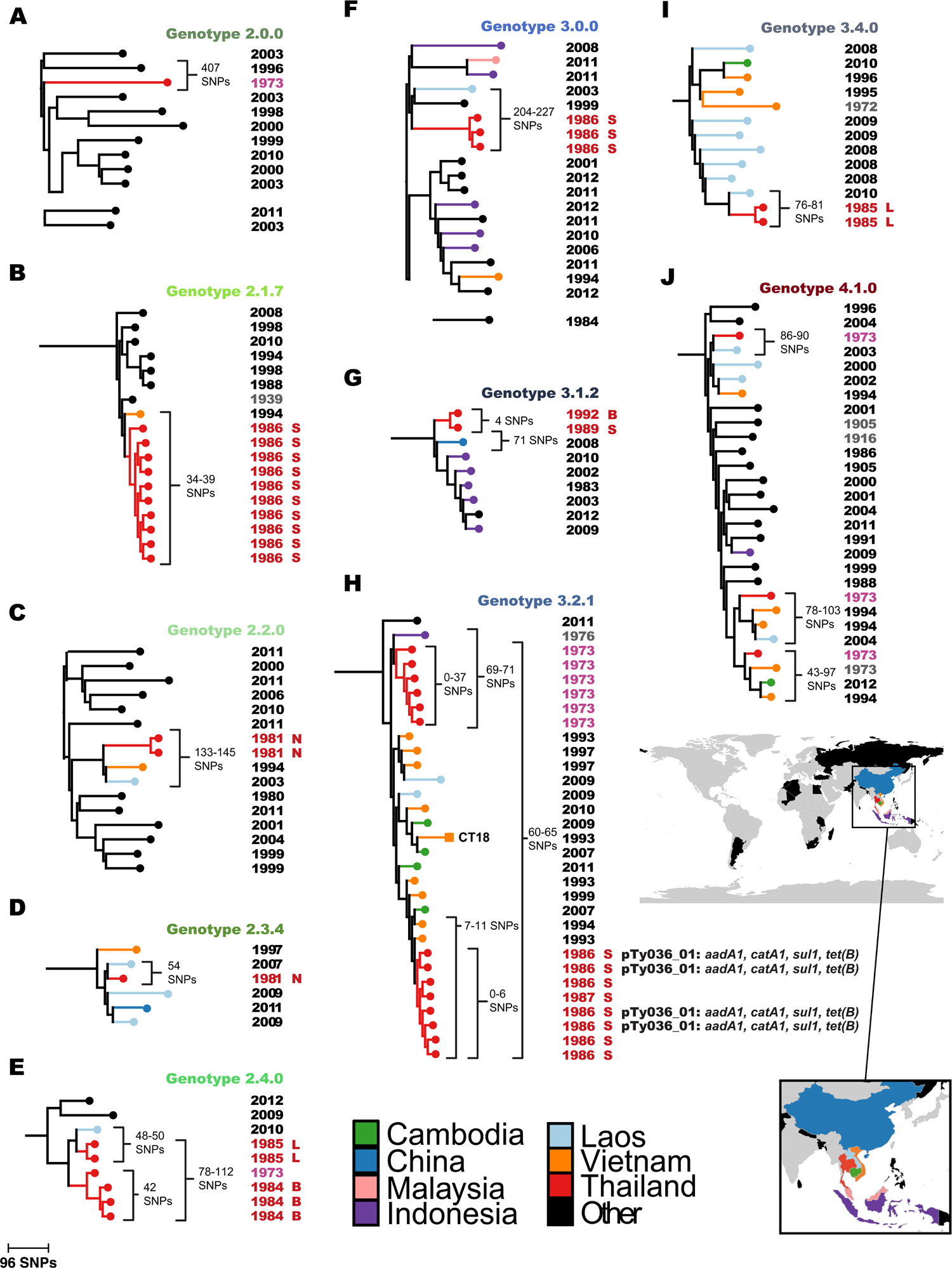
Zoomed in phylogenies showing relationships of Thai S. Typhi to global isolates.

Maximum likelihood trees including S. Typhi isolates from the Thai and global collections are shown, for each genotype that was observed amongst the Thai isolates. Colored branches and nodes indicate country of origin, according to the inset legend. Year of isolation is shown to the left; pink and red, Thai isolates obtained before and after the introduction of the immunization program; grey and black, non-Thai isolates obtained before and after the introduction of the immunization program. Thai isolates are also labelled to indicate their city of origin: L, Loi; B, Bangkok; S, Srakaew; N, Nonthaburi. SNP distances between isolates as well as AMR plasmids are labelled, with any resistance genes indicated in italics. Branch lengths are indicative of the number of SNPs.

**Figure 3.**
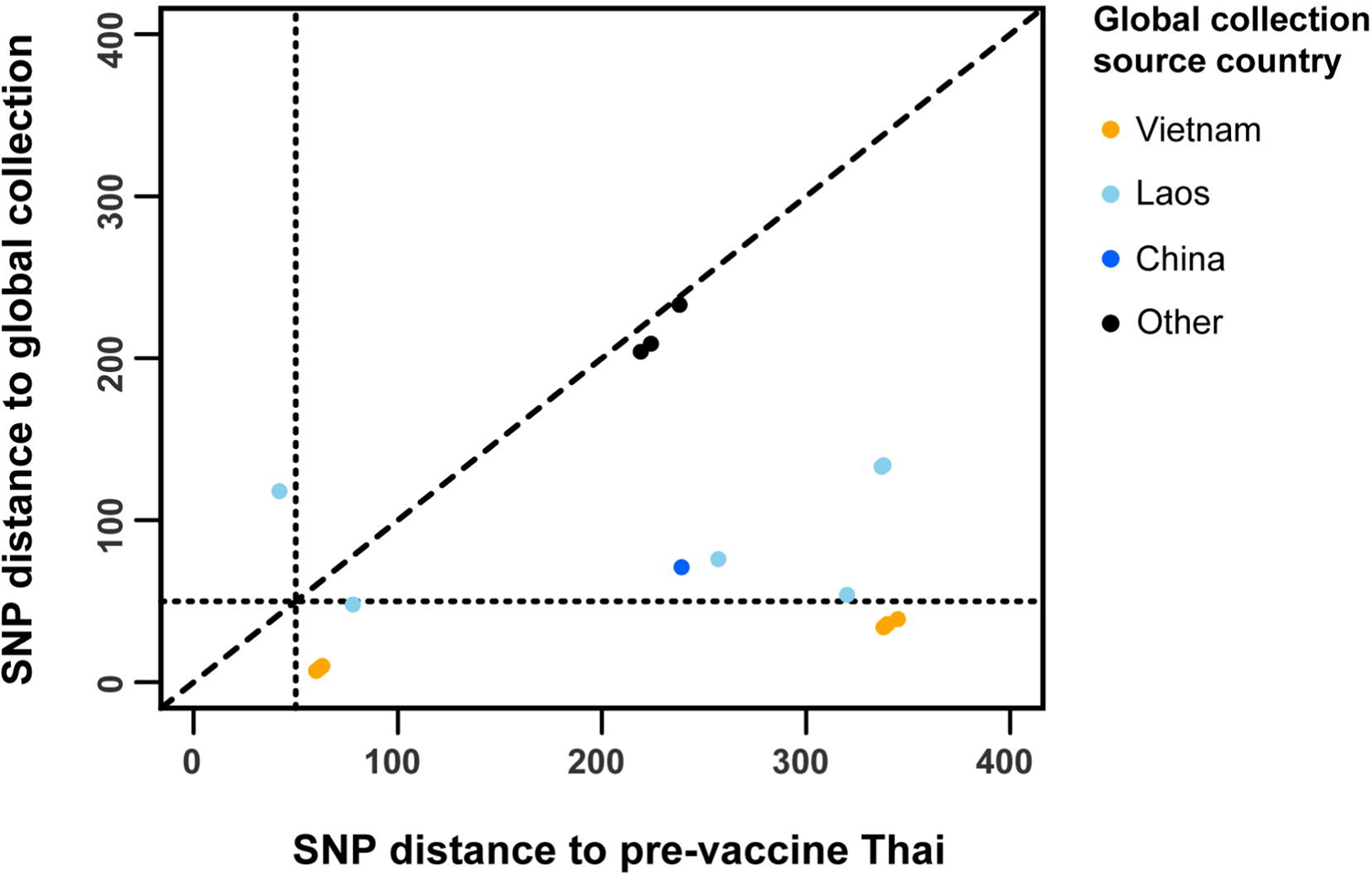
SNP distances for Thai and global collection isolates.

SNP distance between post-vaccine Thai isolates and their closest pre-vaccine Thai and post-vaccine global collection relatives, colored points indicate country of origin.

### Acquired antimicrobial resistance

We identified acquired AMR genes in the genomes of four *S*. Typhi genotype 3.2.1 that were isolated in Srakaew in 1986 **(Fig 1, Table 1).** These isolates shared the same four AMR genes: *sul1*(sulphonamides), *catA1*(chloramphenicol), *tet(B)*(tetracyclines), and *aadA1*(aminoglycosides) which were carried on near-identical plasmids of IncHI1 plasmid sequence type 2 (PST2). Although the presence of insertion sequences (IS) in these plasmids prevented the complete sequences from being assembled, the regions of these plasmids encoding the AMR genes were identical in all assemblies. This commonality suggests they are a single plasmid (referred to as pTy036_01 in **Fig 1** and **Table 1)** that was likely acquired in a common ancestor of this clade. The chromosomal and IncHI1 plasmid sequences for these four isolates were very closely related to those of a 1993 Vietnamese isolate (Viety1-60_1993) in the global *S*. Typhi collection (45), consistent with regional transfer.

### Other plasmids and mobile genetic elements

We identified three non-AMR related plasmids amongst the Thai isolates **(Fig 1, Table 1).** Ty004 (genotype 2.2) carried two novel plasmids that assembled into circular sequences, pTy004_01 and pTy004_02. The largest, pTy004_01, was a novel variant of the cryptic plasmid pHCM2 (29, 49) **(Fig 4).** Ty004 was isolated in Bangkok in 1973, making pTy004_01 the earliest example of a pHCM2-like plasmid reported to date. pTy004_01 was distant from other pHCM2-like plasmids in the global *S*. Typhi genome collection, sharing 92% coverage and 99% nucleotide identity with the reference sequence pHCM2 of *S*. Typhi CT18 (genotype 3.2.1) which was isolated approximately 20 years later in Vietnam (29). The pTy004_01 sequence **(Fig 4)** appears to be ~2 kbp larger than pHCM2, and encodes an additional tRNA-Lys as well as an insertion of a hypothetical protein (*orf17*) into a putative DNA polymerase gene (HCM2.0015c in pHCM2, divided into *orf16* and *orf18* in pTy004_01). Plasmid pTy004_02 was ~38 kbp in size and similar to *E. coli* plasmid pEQ2 (65% coverage, 98% nucleotide identity), encoding genes for conjugation, chromosomal partitioning, addiction systems and an abortive infection protein *(orf44)*. Three isolates (Ty031, Ty042, and Ty049) all of genotype 3.0.0 and obtained from Srakaew in 1986, carried a ~40 kbp cryptic plasmid that we named pTy031_01. This plasmid was similar to that carried by *Enterobacter hormaechei* strain CAV1176 (83% coverage, 96% identity) and encoded genes for chromosomal partitioning, addiction systems, and a putative restriction modification system *(orf33-orf34)*.

**Figure 4.**
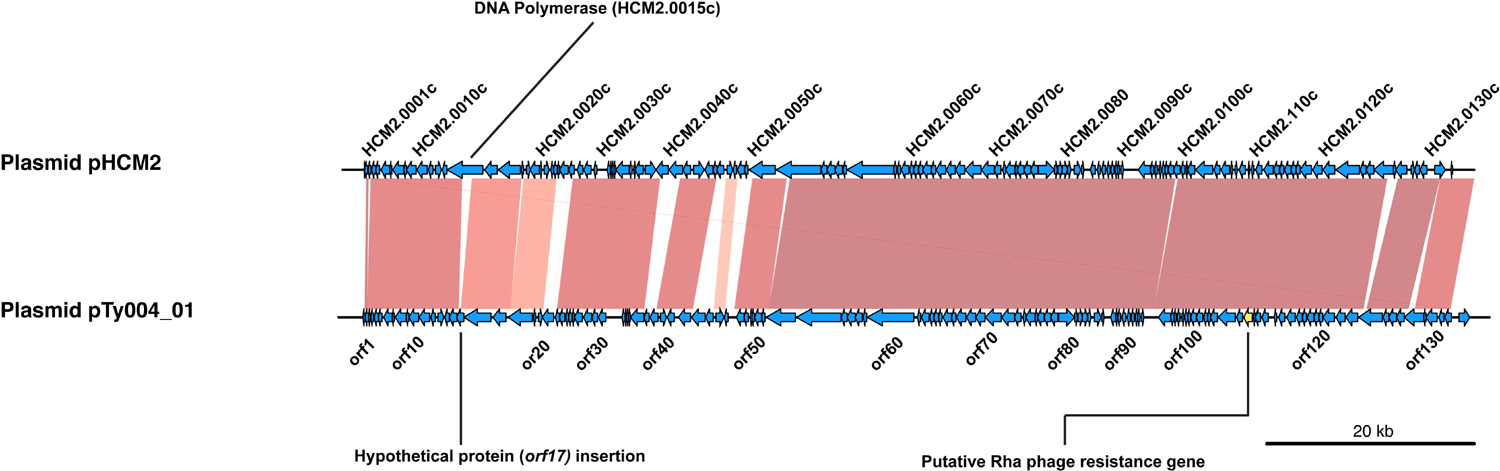
Blast comparison of novel plasmid pTy004_01 with pHCM2 (AL513383).

Shaded regions indicate areas of sequence homology, intensity of shading indicates relative nucleotide similarity. Arrows represent protein coding genes, direction indicates coding strand.

PHAST analysis revealed the presence of novel intact prophages in three Thai *S*. Typhi isolates **(Fig 1, Table 1).** Two *S*. Typhi 3.1.2, isolated from Srakaew in 1989 and Bangkok in 1992, shared a novel phage STYP1 that was similar to fiAA91-ss infective for *Shigella sonnei* **(Fig 5A).** However, the *S*. Typhi phage lacked the cytolethal distending toxin *cdt* genes and the IS2*1* element found in phage fiAA91-ss (50). This prophage sequence had a mosaic architecture, incorporating a number of putative insertions of phage tail fiber genes that were not present in the fiAA91-ss reference genome **(Fig 5A).** Additionally, a single isolate of genotype 4.1 obtained from Bangkok in 1973 contained a novel SfIV-like phage, here named STYP2, that lacked the serotype conversion gene Gtr cluster and IS*1* element of phage SfIV (51). Again, the novel Thai phage variant also encoded novel tail fiber genes not in the SfIV reference genome, as well as a Dam methylase gene *(orf37)***(Fig 5B).**

**Figure 5.**
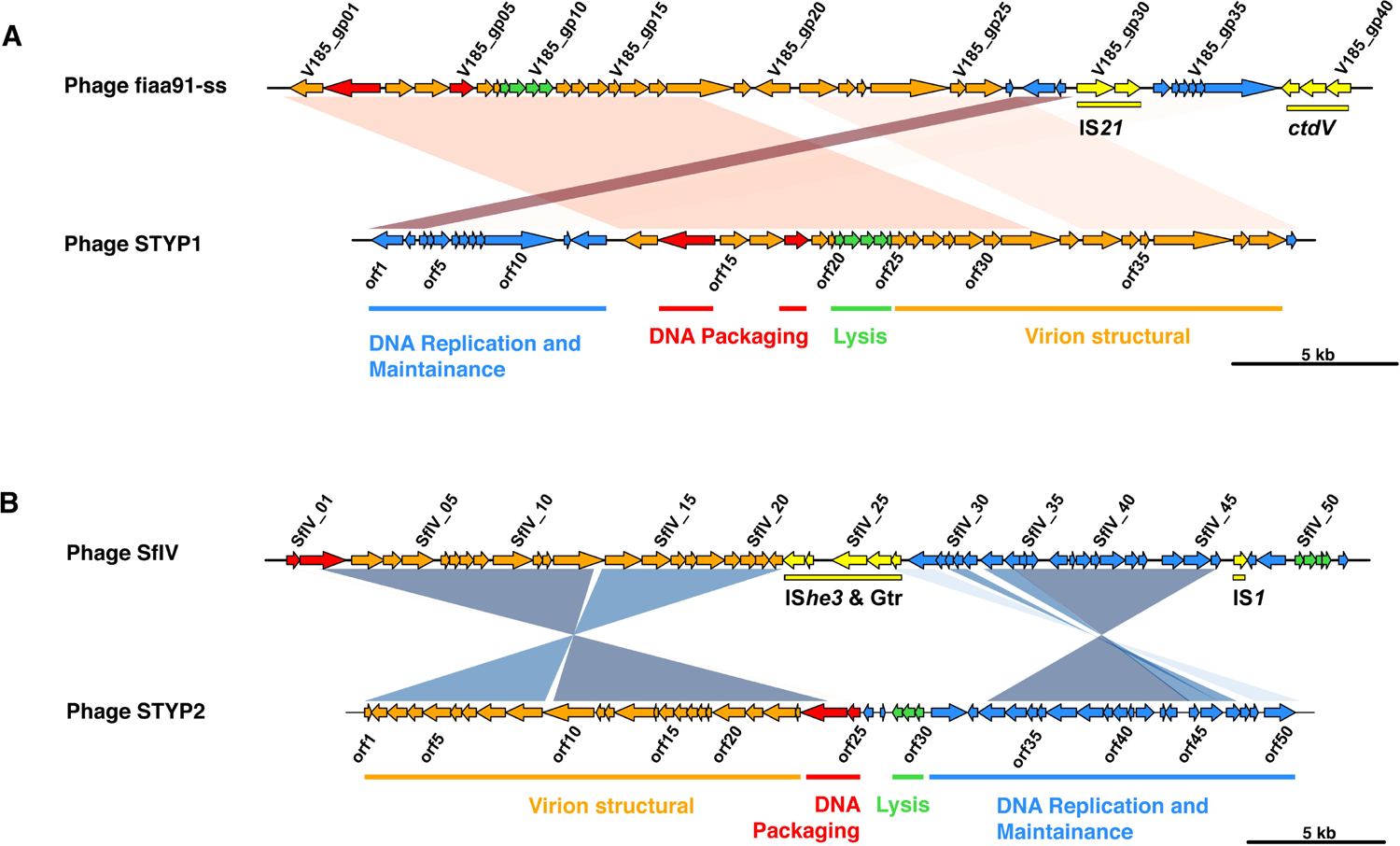
Blast comparison of novel phages observed in Thai S. Typhi isolates to nearest known phage sequences.

(A) Novel phage STYP1 compared to Shigella sonnei phage fiAA91-ss (NC_022750). (B) Novel phage STYP2 compared to Shigella flexneri phage SfIV (NC_022749). Shaded regions indicate areas of sequence homology, intensity of shading indicates relative nucleotide similarity. Arrows represent protein coding genes (direction indicates coding strand), colored by encoded protein functions: red, DNA packaging module; orange, virion morphogenesis module; yellow, cargo genes; blue, DNA replication and lysogenic cycle maintenance; green, lysis module.

## Discussion

These data provide a historical insight into the population structure of *S*. Typhi in Thailand in 1973 (pre-immunization program, n=11) and 1981-1992 (post-immunization program, n=33). It has been reported that the national *S*. Typhi immunization program in Thailand, which commenced in 1977, was highly effective in reducing the burden of typhoid fever (14). Our data are consistent with the hypothesis that the vaccine program successfully depleted the endemic *S*. Typhi population to the extent that most subsequent typhoid cases resulted from sporadic introduction of non-indigenous *S*. Typhi, rather than long-term persistence of the pre-vaccine era population. It is apparent that these introductions were sometimes accompanied by limited local transmissions, resulting in small, localized outbreaks, but we found no evidence to suggest that these result in the establishment of stable local source populations. Notably, the post-immunization *S*. Typhi isolates from Loi (in the north of Thailand near the border with Laos, from which it is separated by the Mekong river) were most closely related to Laos isolates, whilst those from the capital Bangkok and nearby Nonthaburi and Srakaew districts were closely related to other isolates from across Southeast Asia **(Fig 2)**, suggesting there may have been multiple routes of import into Thailand. Our study is limited by the sample of isolates available for analysis, which was small and reflects opportunistic sampling of sporadic local cases in the four sites and historical storage. A larger collection of historical isolates from Thailand and neighboring countries in the 1970s and 1980s would help to further elucidate the epidemiological patterns of *S*. Typhi before and after the vaccination program. However, from our data, it is notable that the Thai isolates cluster according to site, consistent with limited local transmission rather than dissemination of lineages between locations. The only exception to this was two genotype 3.1.2 isolates, which were collected from Srakaew in 1989 and Bangkok in 1992 and differed by only 4 SNPs. This is consistent with either transfer between these cities in Thailand following an initial introduction into the country, or two independent transfers into Thailand from a common source. The phylogenetic structure is most suggestive of the latter, but denser samples from Thailand and/or potential source populations would be required to resolve this with confidence.

While our sample is small, this study is nevertheless the largest to date exploring genetic diversity amongst *S*. Typhi from Thailand. An earlier global haplotyping study that included seven Thai isolates (52) identified five distinct haplotypes in Thailand (H3, 1989; H42, 1990; H50, 2002; Vi-H52, 1990; H79, 2002), three of which are related to genotypes that we identified amongst Thai strains in this study (H79, 2.3.4; H52, 3.4; H42, 3.1.2) (32). Genotype 4.3.1 (H58) was not found amongst our historical Thai isolates. This is consistent with previously published spatiotemporal analyses of the global isolate collection, which showed this rapidly expanding clone only began spreading throughout Asia after 1990 (4). To our knowledge the only evidence to date of the presence of 4.3.1 (H58) in Thailand comes from the global study (4), in which three isolates were identified from 2010-2011, most likely introduced from India. Therefore, our genomic snapshot of the Thai *S*. Typhi population is consistent with previous insights and is likely reasonably representative for the study period. In the years following the vaccination program the prevalence of Typhoid fever in Thailand has continued to decline (53, 54). The vaccination program has been credited with reducing disease incidence in Thailand and was followed by increased economic development in the region as well as improvements to both water and sanitation systems that have likely improved the control of such outbreaks (53, 54). Consequently, Typhoid fever is longer considered a serious public health threat in Thailand (53).

The presence of novel plasmids and prophages in the Thai isolates is also noteworthy. While small plasmids of unknown function have been observed in *S*. Typhi previously (55), they are infrequent compared to the IncHI1 MDR plasmid and the cryptic plasmid pHCM2 (33). Presumably, such plasmids are ephemeral; possibly because their maintenance imposes a fitness burden on the host cells so a strong selective advantage is required for retention (56, 57). It is also possible that the lack of previous reports regarding the diversity of small plasmids in *S*. Typhi reflects a technological complexity, however, this is bypassed with high-throughput WGS and we detected negligible small plasmid content in the global collection of 1,832 genomes using the same screening approach (58). Notably, few of the Thai plasmids share nucleotide sequence homology with those previously described in *S*. Typhi, but were closely related to those found in other *Enterobacteriaceae*. The novel pHCM2-like plasmid (pTy004_01) and two additional plasmids (pTy004_02 and pTy031_01) harbored genes associated with phage resistance, which could provide protection against phage predation (59-62). We also observed two novel prophages integrated into Thai genomes, which both showed variation in their phage tail structural regions compared to close neighbors found in *Shigella/E. coli*. These regions are typically responsible for binding of phage to host receptors (63–65), thus the variation in these regions may be associated with recent adaptations to the *S*. Typhi host. While genomic data from more recent *S*. Typhi collections shows limited evidence for genetic exchange with other organisms (4), the detection amongst older Thai isolates of both phage and plasmids that have been previously associated with *E. coli/Shigella* suggests that genetic exchange may have been more common in the past or in certain localized populations.

Overall, these data provide valuable historical insights into the *S*. Typhi populations circulating in Thailand during the 1970s and 1980s, and early examples of the two most common *S*. Typhi plasmids, as well as other mobile elements identified within the *S*. Typhi population. Importantly, while genomic epidemiology has been applied to study typhoid transmission, antimicrobial resistance evolution and antibiotic treatment failure in various settings (66–68), this study provides an important proof-of-principle demonstration that this approach can also provide useful insights into the impact of typhoid vaccines on circulating bacterial populations. This should motivate the adoption of WGS methods to monitor *S*. Typhi populations during future immunization programs and other large-scale interventions, which could potentially identify differential impacts on distinct genotypes.

## References

1 Parry CM, Hien TT, Dougan G, White NJ, Farrar JJ. Typhoid fever. Ν Engl J Med. 2002;347(22):1770–82.

2 Mogasale V, Maskery B, Ochiai RL, Lee JS, Mogasale VV, Ramani E, et al. Burden of typhoid fever in low-income and middle-income countries: a systematic, literature-based update with risk-factor adjustment. Lancet Glob Health. 2014;2(10):e570–e80.

3 Crump JA, Minte ED. Global trends in typhoid and paratyphoid Fever. Clin Infect Dis. 2010;50(2):241–6.

4 Wong VK, Baker S, Pickard DJ, Parkhill J, Page AJ, Feasey NA, et al. Phylogeographical analysis of the dominant multidrug-resistant H 58 clade of Salmonella Typhi identifies inter-and intracontinental transmission events. Nat Genet 2015;47(6):632–9.

5 Connor BA, Schwartz E. Typhoid and paratyphoid fever in travellers. Lancet Infect Dis. 2005;5(10):623–8.

6 Crump JA, Luby SP, Mintz ED. The global burden of typhoid fever. Bull World Health Organ. 2004;82(5):346–53.

7 Sur D, von Seidlein L, Manna B, Dutta S, Deb AK, Sarkar BL, et al. The malaria and typhoid fever burden in the slums of Kolkata, India: data from a prospective community-based study. Trans R Soc Trop Med Hyg. 2006;100(8):725–33.

8 Karkey A, Arjyal A, Anders KL, Boni MF, Dongol S, Koirala S, et al. The Burden and Characteristics of Enteric Fever at a Healthcare Facility in a Densely Populated Area ofKathmandu. PLoS ONE. 2010;5(ll):el3988.

9 Cairncross S, Cumming O, Jeandron A, Rheingans R, Ensink J, Brown J. Water, Sanitation and Hygiene Evidence Paper. London: Department for International Development. 2013.

10 Bhutta ZA. Current concepts in the diagnosis and treatment of typhoid fever. BMJ. 2006;333(7558):78–82.

11 Chau TT, Campbell JI, Galindo CM, Van Minh Hoang N, Diep TS, Nga TTT, et al. Antimicrobial Drug Resistance of Salmonella enterica Serovar Typhi in Asia and Molecular Mechanism of Reduced Susceptibility to the Fluoroquinolones. Antimicrob Agents Chemother. 2007;51(12):4315–23.

12 Kariuki S, Revathi G, Muyodi J, Mwituria J, Munyalo A, Mirza S, et al. Characterization of Multidrug-Resistant Typhoid Outbreaks in Kenya. J Clin Microbiol. 2004;42(4): 1477–82.

13 Holt KE, Phan MD, Baker S, Duy PT, Nga TVT, Nair S, et al. Emergence of a Globally Dominant IncHIl Plasmid Type Associated with Multiple Drug Resistant Typhoid. PLoS Negl Trop Dis. 2011;5(7):el245.

14 Bodhidatta L, Taylor DN, Thisyakom U, Echeverria P. Control of Typhoid Fever in Bangkok, Thailand, by Annual Immunization of Schoolchildren with Parenteral Typhoid Vaccine. Rev Infect Dis. 1987;9(4):841–5.

15 Klugman K, Koornhof H, Schneerson R, Cadoz M, Gilbertson I, Robbins J, et al. Protective activity of Vi capsular polysaccharide vaccine against typhoid fever. Lancet. 1987;330(8569):1165–9.

16 Levine M, Black R, Ferreccio C, Germanier R. Large-scale field trial of Ty21A live oral typhoid vaccine in enteric-coated capsule formation. Lancet 1987;329(8541):1049–52.

17 Ashcroft MT, Singh Β, Nicholson CC, Ritchie JM, Sobryan E, Williams F. A seven-year field trial of two typhoid vaccines in Guyana. Lancet 1967;290(7525):1056–9.

18 Waddington CS, Darton TC, Jones C, Haworth K, Peters A, John T, et al. An Outpatient, Ambulant-Design, Controlled Human Infection Model Using Escalating Doses of Salmonella Typhi Challenge Delivered in Sodium Bicarbonate Solution. Clin Infect Dis. 2014;58(9):1230–40.

19 Scobie HM, Nilles E, Kama M, Kool JL, Minte E, Wannemuehler ΚΑ, et al. Impact of a targeted typhoid vaccination campaign following cyclone Tomas, Republic of Fiji, 2010. Am J Trop Med Hyg. 2014;90(6):1031–8.

20 Thompson CN, Kama M, Acharya S, Bera U, Clemens J, Crump JA, et al. Typhoid fever in Fiji: a reversible plague? Trop Med Int Health. 2014;19(10):1284–92.

21 Whitaker JA, Franco-Paredes C, del Rio C, Edupuganti S. Rethinking typhoid fever vaccines: implications for travelers and people living in highly endemic areas. J Travel Med. 2009;16(1):46–52.

22 DeRoeck D, Ochiai RL, Yang J, Anh DD, Alag V, Clemens JD. Typhoid vaccination: the Asian experience. Expert Rev Vaccines. 2008;7(5):547–60.

23 Guzman CA, Borsuteky S, Griot-Wenk M, Metcalfe IC, Pearman J, Collioud A, et al. Vaccines against typhoid fever. Vaccine. 2006;24(18):3804–11.

24 Ochiai RL, Acosta CJ, Agtini M, Bhattacharya SK, Bhutta ZA, Do CG, et al. The Use of Typhoid Vaccines in Asia: The DOMI Experience. Clin Infect Dis. 2007;45(Supplement 1):S34–S8.

25 CLSI. Performance standards for anitmicrobial disk suseptibility tests. Approved standard. CLSI document M02. Wayne, Pa: CLSI; 1984.

26 CLSI. Performace standards for antimicrobial disk susceptibility tests. Tentative standard. CLSI document M02. Wayne, Pa: CLSI; 1979.

27 CLSI. Performance standards of antimicrobial disk susceptibility tests. Tentative standard. = CLSI document M02-A4. Wayne, Pa: CLSI; 1988.

28 CLSI. Performace standards for anitmicrobial disk susceptibility tests. Approved Standard. CLSI document = M02-A4. Wayne, Pa: CLSI; 1990.

29 Parkhill J, Dougan G, James KD, Thomson NR, Pickard D, Wain J, et al. Complete genome sequence of a multiple drug resistant Salmonella enterica serovar Typhi CT18. Nature. 2001;413(6858):848–52.

30 Langmead Β, Salzberg SL. Fast gapped-read alignment with Bowtie 2. Nat Meth. 2012;9(4):357–9.

31 Li H, Handsaker B, Wysoker A, Fennell T, Ruan J, Homer Ν. The sequence alignment/Map format and SAMtools. Bioinformatics. 2009;25.

32 Wong VK, Baker S, Connor TR, Pickard D, Page AJ, Dave J, et al. An extended genotyping framework for Salmonella enterica serovar Typhi, the cause of human typhoid. Nat Commun. 2016;7:12827.

33 Holt KE, Parkhill J, Mazzoni CJ, Roumagnac P, Weill F-X, Goodhead I. High-throughput sequencing provides insights into genome variation and evolution in Salmonella Typhi. Nat Genet 2008;40.

34 Croucher NJ, Page AJ, Connor TR, Delaney AJ, Keane JA, Bentley SD, et al. Rapid phylogenetic analysis of large samples of recombinant bacterial whole genome sequences using Gubbins. Nucleic Acids Res. 2014.

35 Holt KE, Thomson NR, Wain J, Langridge GC, Hasan R, Bhutta ZA, et al. Pseudogene accumulation in the evolutionary histories of Salmonella enterica serovars Paratyphi A and Typhi. BMC genomics. 2009;10:36.

36 Stamatakis A. RAxML-VI-HPC: maximum likelihood-based phylogenetic analyses with thousands of taxa and mixed models. Bioinformatics. 2006;22(21):2688–90.

37 Paradis E, Claude J, Strimmer K. APE: Analyses of Phylogenetics and Evolution in R language. Bioinformatics. 2004;20(2):289–90.

38 Gupta SK, Padmanabhan BR, Diene SM, Lopez-Rojas R, Kempf M, Landraud L, et al. ARG-ANNOT, a New Bioinformatic Tool To Discover Antibiotic Resistance Genes in Bacterial Genomes. Antimicrob Agents and Chemother. 2014;58(1):212–20.

39 Inouye M, Dashnow H, Raven L-A, Schultz MB, Pope BJ, Tomita T. SRST2: rapid genomic surveillance for public health and hospital microbiology labs. Genome Med. 2014;6.

40 Carattoli A, Bertini A, Villa L, Falbo V, Hopkins KL, Threlfall EJ. Identification of plasmids by PCR-based replicon typing. J Microbiol Meth. 2005;63(3):219–28.

41 Carattoli A, Zankari E, Garcia-Fernândez A, Voldby Larsen M, Lund O, Villa L, et al. In Silico Detection and Typing of Plasmids using PlasmidFinder and Plasmid Multilocus Sequence Typing. Antimicrob Agents Chemother. 2014;58(7):3895–903.

42 Bankevich A, Nurk S, Antipov D, Gurevich AA, Dvorkin M, Kulikov AS, et al. SPAdes: A New Genome Assembly Algorithm and Its Applications to Single-Cell Sequencing. J Comp Biol. 2012;19(5):455–77.

43 Wick RR, Schultz MB, Zobel J, Holt KE. Bandage: interactive visualisation of de novo genome assemblies. Bioinformatics. 2015.

44 Seemann T. Prokka: rapid prokaryotic genome annotation. Bioinformatics. 2014.

45 Phan M-D, Kidgell C, Nair S, Holt KE, Turner AK, Hinds J, et al. Variation in Salmonella enterica Serovar Typhi IncHI1 Plasmids during the Global Spread of Resistant Typhoid Fever. Antimicrob Agents Chemother. 2009;53(2):716–27.

46 Jolley K, Maiden M. BIGSdb: Scalable analysis of bacterial genome variation at the population level. BMC Bioinformatics. 2010;11(1):595.

47 Zhou Y, Liang Y, Lynch KH, Dennis JJ, Wishart DS. PHAST: a fast phage search tool. Nucleic Acids Res. 2011;39(Web Server issue):W347–52.

48 Guy L, Roat Kultima J, Andersson SGE. genoPlotR: comparative gene and genome visualization in R. Bioinformatics. 2010;26(18):2334–5.

49 Kidgell C, Pickard D, Wain J, James K, Diem Nga LT, Diep TS, et al. Characterisation and distribution of a cryptic Salmonella typhi plasmid pHCM2. Plasmid. 2002;47(3):159–71.

50 Allue-Guardia A, Imamovic L, Muniesa M. Evolution of a self-inducible cytolethal distending toxin type V-encoding bacteriophage from Escherichia coli 0157:H7 to Shigella sonnei. J Virol. 2013;87(24):13665–75.

51 Jakhetia R, Talukder K, Verma N. Isolation, characterization and comparative genomics of bacteriophage SfIV: a novel serotype converting phage from Shigella flexneri. BMC genomics. 2013;14(1):677.

52 Roumagnac P, Weill FX, Dolecek C, Baker S, Brisse S, Chinh NT, et al. Evolutionary history of Salmonella typhi. Science. 2006;314.

53 DeRoeck D, Clemens JD, Nyamete A, Mahoney RT. Policymakers' views regarding the introduction of new-generation vaccines against typhoid fever, shigellosis and cholera in Asia. Vaccine. 2005;23(21):2762–74.

54 IHME. Global Burden of Disease Study 2015 Seattle, USA: Institute for Health Metrics and Evaluation; 2015 [cited 2016 08/12/2016]. Available from: http://ghdx.healthdata.org/gbd-results-tool.

55 Holt KE, Perkins TT, Dougan G, Kingsley RA. Genomics and pathogenesis of Salmonella enterica serovars Typhi and Paratyphi A. Salmonella: From genome to function. 2011:107–21.

56 Rychlik I, Gregorova D, Hradecka H. Distribution and function of plasmids in Salmonella enterica. Vet Microbiol. 2006;112(1):1–10.

57 Murray BE, Levine MM, Cordano AM, D'Ottone Κ, Jayanetra Ρ, Kopecko D, et al. Survey of plasmids in Salmonella typhi from Chile and Thailand. J Infect Dis. 1985;151(3):551–5.

58 jørgensen TS, Xu Ζ, Hansen MA, Sørensen SJ, Hansen LH. Hundreds of Circular Novel Plasmids and DNA Elements Identified in a Rat Cecum Metamobilome. PLoS ONE. 2014;9(2):e87924.

59 Buckling A, Rainey PB. Antagonistic coevolution between a bacterium and a bacteriophage. Ρ Roy Soc B-Biol Sci. 2002;269(1494):931–6.

60 Koskella Β, Brockhurst MA. Bacteria-phage coevolution as a driver of ecological and evolutionary processes in microbial communities. FEMS microbiology reviews. 2014;38(5):916–31.

61 Lenski R. Coevolution of Bacteria and Phage: Are There Endless Cycles of Bacterial Defenses and Phage Counterdefenses? J Theor Biol. 1984;108:319–25.

62 Schräg SJ, Mittler JE. Host-parasite coexistence: The role of spatial refuges in stabilizing bacteria-phage interactions. Am Nat. 1996;148(2):348–77.

63 Kutter E, Raya R, Carlson K. Molecular mechanisms of phage infection. In: Kutter E, Sulakvelidze A, editors. Bacteriophages Biology and Applications. Florida: CRC Press; 2005.

64 Guttman B, Raya R, Kutter E. Chapter 3-Basic Phage Biology. In: Kutter E, Sulakvelidze A, editors. Bacteriophages-Biology and Applications. New York: CRC Press; 2005. p. 29–66.

65 Thomson N, Baker S, Pickard D, Fookes M, Anjum M, Hamlin N, et al. The role of prophage-like elements in the diversity of Salmonella enterica serovars. J Mol Biol. 2004;339(2):279–300.

66 International Typhoid C, Wong VK, Holt KE, Okoro C, Baker S, Pickard DJ, et al. Molecular Surveillance Identifies Multiple Transmissions of Typhoid in West Africa. PLOS Negl Trop Dis. 2016;10(9):e0004781.

67 Pham Thanh D, Thompson CN, Rabaa MA, Sona S, Sopheary S, Kumar V, et al. The Molecular and Spatial Epidemiology of Typhoid Fever in Rural Cambodia. PLoS Negl Trop Dis. 2016;10(6):e0004785.

68 Pham Thanh D, Karkey A, Dongol S, Ho Thi N, Thompson CN, Rabaa MA, et al. A novel ciprofloxacin-resistant subclade of H58 Salmonella Typhi is associated with fluoroquinolone treatment failure. eLife. 2016;5:el4003.

